# Extracellular matrix proteins modulate lymphatic endothelial cell junction morphology and barrier function

**DOI:** 10.64898/2026.01.30.702887

**Authors:** Sarfaraz Ahmad Ejazi, Aisha Abdulkarimu, Lina Berhaneyessus, Andrej Radoja, Katharina Maisel

**Affiliations:** Fischell Department of Bioengineering, University of Maryland, College Park

**Author notes:** Correspondence to: Dr. Katharina Maisel, Fischell Department of Bioengineering University of Maryland, College Park.

## Abstract

The extracellular matrix (ECM) plays a pivotal role in lymphatic vasculature physiology, yet the specific contribution of individual ECM components to lymphatic endothelial permeability remains poorly understood, limiting the development of physiologically relevant in vitro models for lymphatic disease research and therapeutic development. Here, we used an in vitro transwell platform to systematically investigate how four clinically relevant ECM proteins, collagen I, fibronectin, fibrin, and laminin, regulate human lymphatic endothelial cell (LEC) barrier function and junctional integrity. Fibrin and collagen I substrates enhanced barrier integrity, demonstrating 80% and 67% increases in transendothelial electrical resistance (TEER), respectively, compared to uncoated controls. FITC-dextran transport assays confirmed these findings, with fibrin and collagen I reducing permeability by 20% and 10%, respectively. Immunofluorescence analysis revealed elevated ZO-1 expression on fibrin, fibronectin, and laminin matrices, while VE-cadherin levels remained unchanged across conditions. Quantitative junctional analysis demonstrated that fibrin increased ZO-1 junction continuity by ∼35%, while collagen I and fibronectin enhanced continuity by ∼22%, with all ECM coatings reducing discontinuous junctions by 60–80%. Mechanistically, RhoA expression was reduced in LECs cultured on fibrin, suggesting decreased stress fiber formation contributes to enhanced barrier function, though overall actin cytoskeletal anisotropy remained unchanged. These findings demonstrate that ECM composition modulates LEC junctional organization and barrier integrity, with fibrin and collagen I exerting the most pronounced barrier-enhancing effects. This engineered platform provides a foundation for developing next-generation in vitro models of lymphatic vasculature that more accurately recapitulate physiological conditions, with applications in lymphedema research, cancer metastasis studies, and immune cell trafficking investigations.

## INTRODUCTION

Lymphatic vessels work in conjunction with the blood vascular system to function as a unidirectional conduit for continuous filtration and removal of fluid from interstitial tissue through lymph nodes and back to systemic circulation ^1,2^. The lymphatic vasculature forms a conduit of polarized lymphatic endothelial cells (LECs), surrounded by connective tissues and matrix proteins that together form the interstitial tissue space ^3^. The structural component of this interstitial space is the extracellular matrix (ECM) and basement membrane (BM), a specialized ECM that envelops the lymphatic vasculature with discontinuity in initial lymphatics and sleeve-like coverage in collecting lymphatics. The initial lymphatic vasculature is characterized by its increased permeability, allowing fluids, cells, proteins, and other materials to transport from peripheral tissues into lymphatic vessels. Collecting lymphatic vessels are more similar to veins, with valves and smooth muscles that facilitate propelling of fluid from tissue to the lymph nodes and into systemic circulation. The ECM and BM provide structural support to both vessels and also allow for transmission of biochemical signals to cells via specific cell surface receptors, such as integrins or anchoring filaments ^4–7^. The passage of solute and fluid across LECs and into the vessel is regulated by the tight and adherens cell-cell junctions ^8^. It has been postulated that the BM may also significantly contribute to this, given its incomplete nature in lymphatic initial vessels that allows LECs to anchor to the interstitial ECM and thus permits the uptake of solute and fluid into the vessel lumen^7^.

The ECM is composed of key structural proteins (i.e., collagen I, III, IV), glycoproteins (such as fibronectin), and glycosaminoglycans such as hyaluronan that provide structural and functional support to lymphatics^7,9^. The BM is composed primarily of collagen IV/ XVIII, laminin, fibronectin, structural linkers, the glycoproteins nidogen-1 and entactin, as well as heparan sulfate proteoglycans ^7^. These components are produced by LECs and supporting mural cells (pericytes or smooth muscle cells) and ultimately required for the maintenance and integrity of LV; without them LV function is compromised ^7,10^. Among these, collagen, fibronectin, and laminin play a crucial role in providing a structural scaffold and anchoring cells to the BM.

Collagen is a key protein that supports and maintains the structure of the lymphatic vasculature ^9,11^. LECs bind to collagen within the ECM via integrins α1β1 and α2β1 ^9,11^. Previous studies have shown that collagen type I specifically enhances LEC survival, maintains LEC spindle morphology, and supports a tubular, capillary-like structure common to vascular endothelial cells, as observed *in vivo* ^11–14^. This suggests that collagens are critical for maintaining LEC morphology and the vessel integrity.

Fibronectin, a glycoprotein, facilitates anchoring and adhering LECs to the ECM through integrin α5β1^9^. LECs have better proliferation and adhesion and can form tube-like formations when cultured on fibronectin *in vitro* ^12,13^. More recently, researchers maintained LECs on fibronectin-coated surfaces *in vitro* to maximize cell adhesion and proliferation ^14^. This suggests that fibronectin is important for maintaining LEC proliferation and plasticity.

Laminin, a large trimeric glycoprotein and a major component of the BM, adhesion to LECs is facilitated β1 integrins^4,15,16^. LECs mainly express laminin isoforms 411 (⍺4β1𝛾𝛾1) and 511(⍺5β1𝛾𝛾1), which are crucial for BM assembly and maintenance ^10,17^. Laminin has been shown to regulate VE-cadherin localization in postcapillary venules ^15^. Researchers observed a decrease in VE-cadherin expression in mouse skin tissue in laminin ⍺4 (lama 4-/-) and laminin ⍺5 (lama 5-/-) knockout mice, compared to the untreated control ^15^. This suggests that laminin is critical for maintaining VE-cadherin expression on LECs, as well as other lymphatic markers such as LYVE-1 ^10^.

Recently, lymphatic researchers have utilized fibrin hydrogel to investigate the underlying mechanisms regulating lymphangiogenesis, due to fibrin’s roles in wound healing and tissue repair ^18,19^. Fibrin is a fibrous protein formed from fibrinogen in the presence of thrombin during the blood clotting process. It is considered abundant during this process and at the site of injuries. In the presence of fibrin, LECs form capillary-like, tubular structures, indicative of functional lymphatic vasculature ^19–23^. This suggests that fibrin is also critical for lymphatic tube formation and vessel integrity.

We sought to systematically investigate how LECs interact with different components of the ECM specifically how they affect LEC barrier function and cell-cell junction integrity, given the majority of the work in the field focused on the effect of ECM on specific lymphatic markers, tube formation, or lymphangiogenesis. We cultured LECs on collagen, fibronectin, laminin, and fibrin-coated transwells and explored how each substrate affects cell-cell junction morphology and lymphatic permeability. Fibrin induced a more continuous LEC junctional architecture and suppressed punctate, perpendicular ZO-1 morphology. Fibrin matrix also promotes a less permeable, more robust cell-cell junctions, and this may be mediated through an interaction between the ECM and the actin cytoskeleton, initiated by RhoA. Our findings illustrate the direct implications the ECM has on the architecture of both adherens and tight junctions and ultimately the lymphatic barrier, making ECM a critical parameter for studying LECs in vitro and engineering complex models of lymphatic vasculature.

## RESULTS

### ECM proteins modulate lymphatic permeability

We found that fibrin and collagen increased LEC barrier properties as indicated by increased transendothelial electrical resistance (TEER) and decreased transport of a fluorescently labeled low molecular weight dextran. Human LECs were seeded on transwells coated with collagen, fibronectin, laminin, and fibrin along with PBS as no-ECM control (**Figure 1A**). LECs were cultured for 24, 48, and 72 hours prior to immunostaining. Cell count and immunostaining of ZO-1 and VE-cadherin were quantified, showing no significant difference across the different ECM coatings up to 72 hours, before a complete monolayer was formed, indicating these ECM proteins do not directly affect LEC proliferation (**Supplementary Figure 1A**). Upon reaching confluence, typically within 5–7 days, LEC monolayers on each ECM protein were evaluated for barrier integrity. Transendothelial electrical resistance (TEER) demonstrated that fibrin and collagen had the significantly highest values (36 ± 2 and 34 ± 2 ohms.cm^2^, respectively), followed by fibronectin (26 ± 2 ohms.cm^2^), compared to the no-ECM, PBS control (20 ± 3 ohms.cm^2^) (**Figure 1B**). Laminin-coated monolayers, in contrast, caused a reduction (12 ± 3 ohms.cm^2^) in TEER compared to the control.

**Figure 1.**
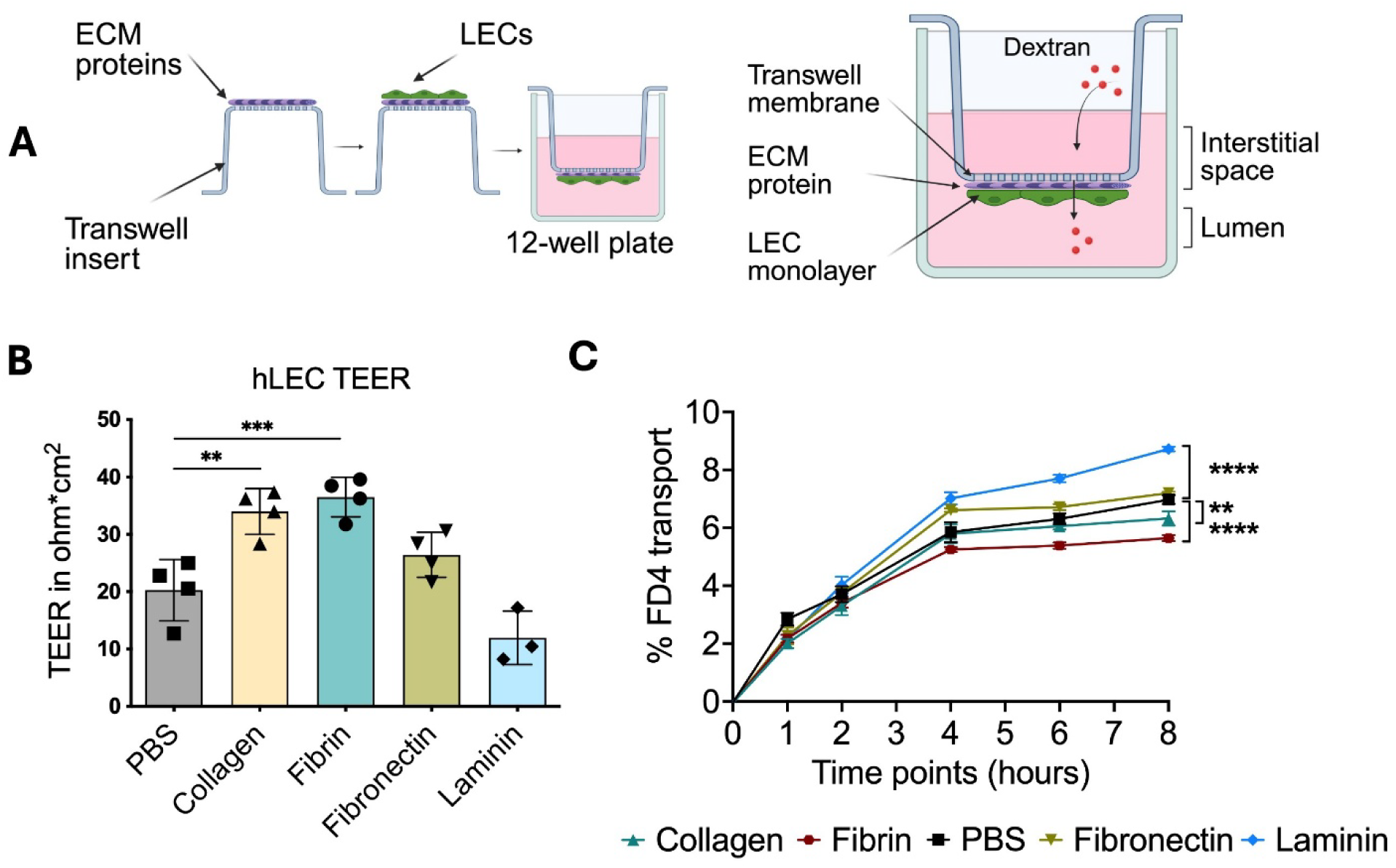
Collagen and fibrin promote hLEC monolayer integrity; where laminin permeabilizes. **(A)** Schematic of *in vitro* transport assay with different ECM coatings on transwells. **(B)** TEER measurements of hLEC monolayers with different ECM coatings compared to no-ECM, PBS control. Multiple comparison One-way Anova as performed with post-hoc Dunnett test for TEER comparison with PBS **(C)** Transport of FITC-Dextran (4 kDa) across hLECs monolayer with different ECM proteins over 8 hours. Data represented as mean ± SEM from n=4 experiments. Two-Way Anova was performed with post-hoc Dunnett test for comparison where significance levels are indicated as * p < 0.05, ** p < 0.01, *** p < 0.001, **** p < 0.0001; ns, not significant.

LEC monolayer permeability was further assessed using FITC-dextran (∼4kDA; FD4), a bio-inert sugar molecule capable of crossing between cells. Consistent with TEER results, dextran permeability was the lowest across LECs grown on fibrin- and collagen with 5.7% and 6.4% FD4 transport across the monolayer after 8 hours, respectively, compared to 7.0% in the PBS control. However, fibronectin caused no significant difference with 7.2% FD4 transport, and laminin displayed a substantial increase in permeability with 8.7% transport (**Figure 1C**).

**Supplementary Figure 1.**
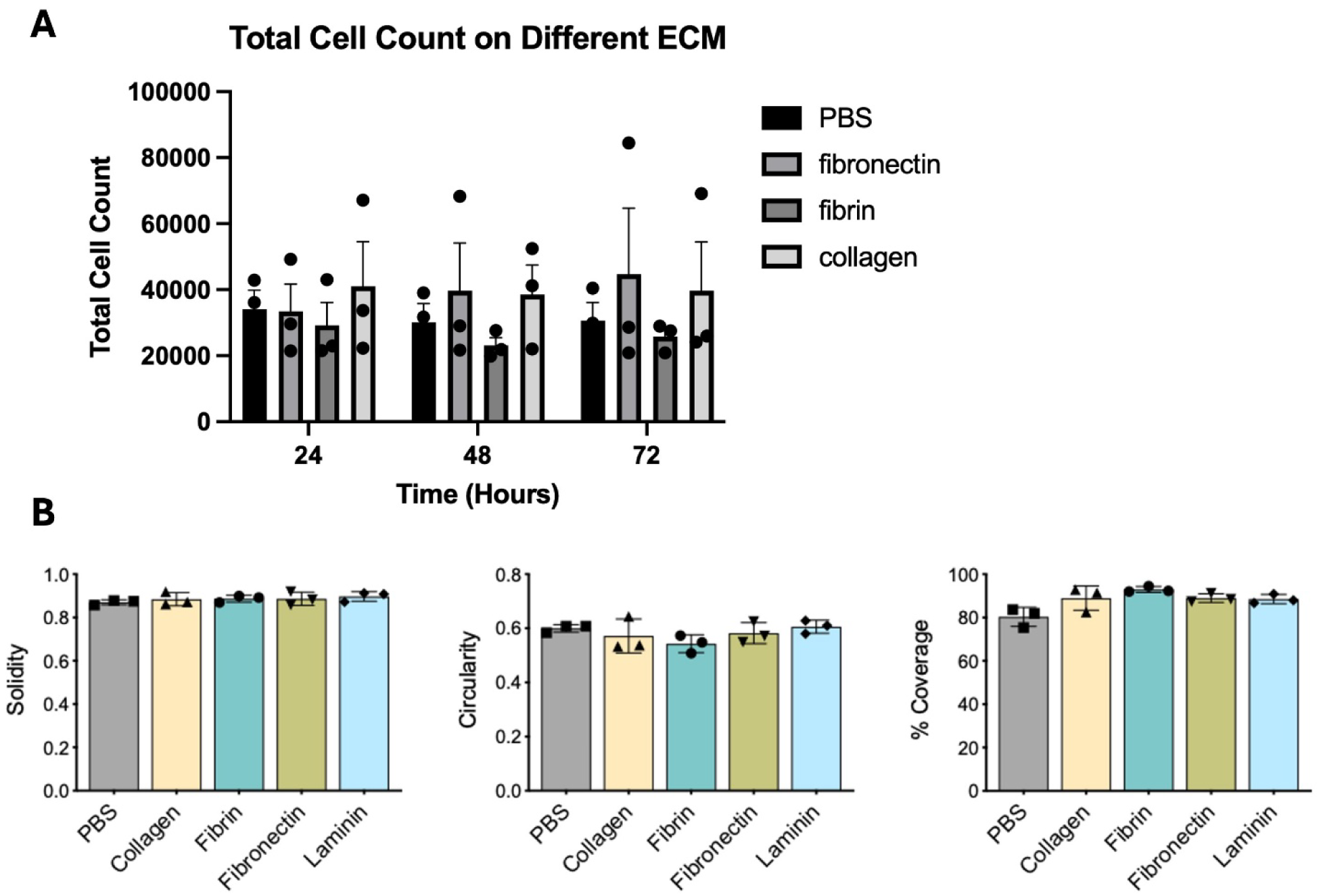
ECM coating does not affect hLEC proliferation and morphology. (A) No significant differences in cell counts were seen in ECM proteins fibronectin, fibrin, and collagen compared to PBS control over 72 hours. **(B)** No significant difference in solidity, circularity, and percent junction protein coverage of VE-cadherin along cell perimeter between different ECM substrates. Data are presented as mean ± SEM from n <2 experiments. One-way ANOVA and unpaired t-test were performed where significance levels are indicated as * p < 0.05, ** p < 0.01, *** p < 0.001, **** p < 0.0001; ns, not significant

### ECM affects the morphology of lymphatic cell-cell junctions

We found that fibrin increased the continuity of the tight junction protein ZO-1 and caused an increase in its expression, suggesting this may be in part the mechanism for changes in permeability observed with TEER and dextran transport. Tight junction protein ZO-1 and adherens junction protein VE-cadherin are key regulators of lymphatic barrier function. Immunofluorescence imaging revealed visibly enhanced expression of both proteins in LECs cultured on fibrin and collagen compared to the PBS control (**Figure 2A**). The mean fluorescence intensity (MFI) analysis revealed a significant 1.2-fold increase in ZO-1 expression in LEC cultured on fibrin (**Figure 2B**) compared to no-ECM control. In contrast, VE-cadherin MFI remained unchanged. Additionally, F-actin staining demonstrated no appreciable difference in stress fiber formation, as anisotropy values were below 1 across all conditions, indicating randomly oriented actin fibers (**Figure 2C**).

**Figure 2.**
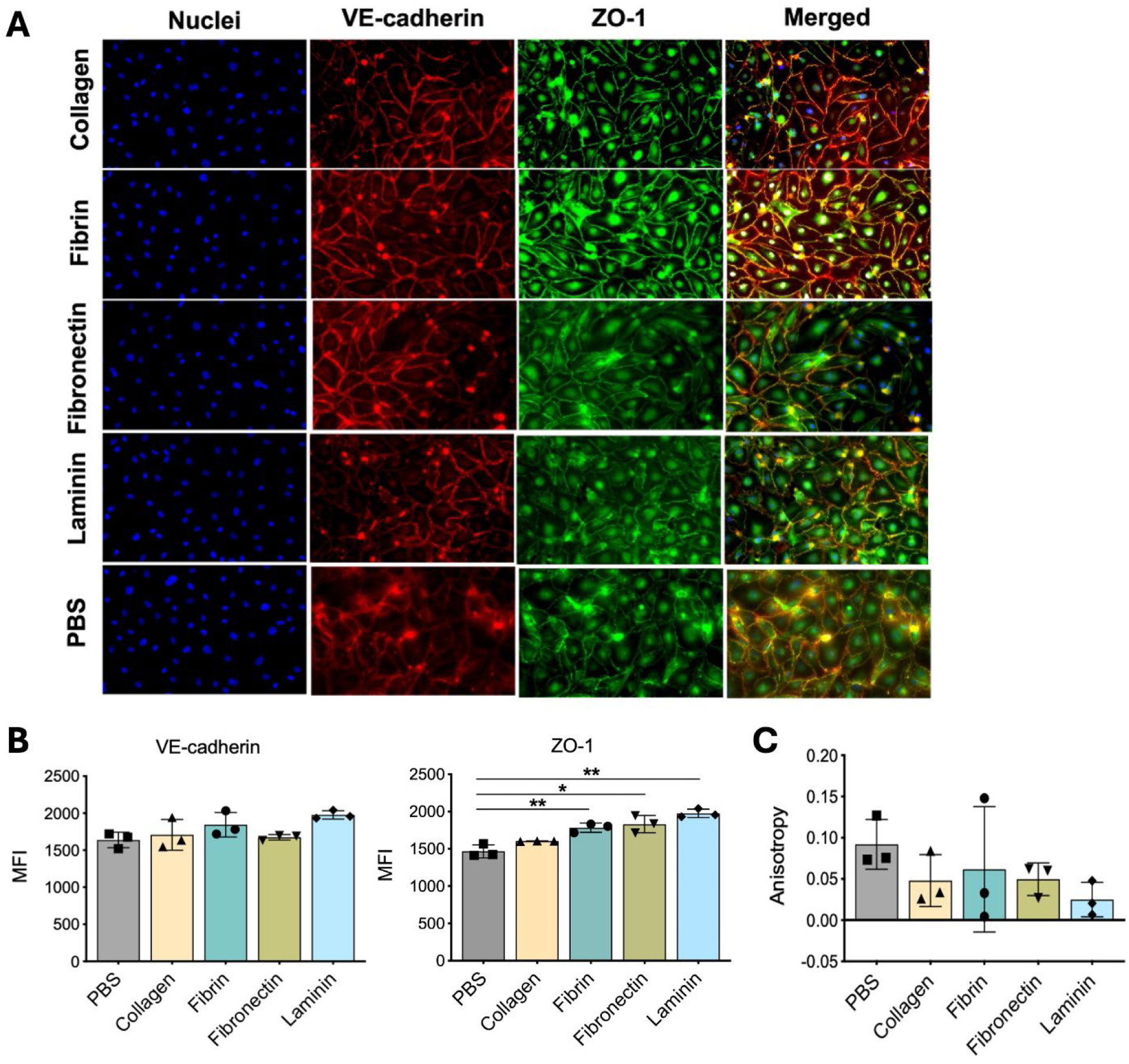
Immunofluorescence imaging displays comparable junction morphology across ECM coatings. (A) Representative immunofluorescence images of VE-cadherin and ZO-1 expression on different ECM coatings. (B) Quantified Mean Fluorescence Intensity (MFI) of VE-cadherin and ZO-1 expression. (C ) Qiamtofoed stress fiber alignment of F-actin. Nonparametric Welch’s test was performed for statistical difference. Data presented as mean ± SEM from n=3 representative images for each groups. Welch’s t-test was performed for statistical analysis where significance levels are indicated as * p < 0.05, ** p < 0.01, *** p < 0.001, **** p < 0.0001; ns, not significant.:

To quantify further changes in LECs’ morphological differences when cultured on different ECM proteins, we used the Junction Analyzer Program (JAnaP) to characterize the immunostaining of LEC junction, tight and adherens junction protein (**Figure 3A**). ZO-1 analysis showed minimal differences in cell solidity and circularity, but a modest increase in monolayer junctional coverage (**Supplementary Figure 1B**).

**Figure 3.**
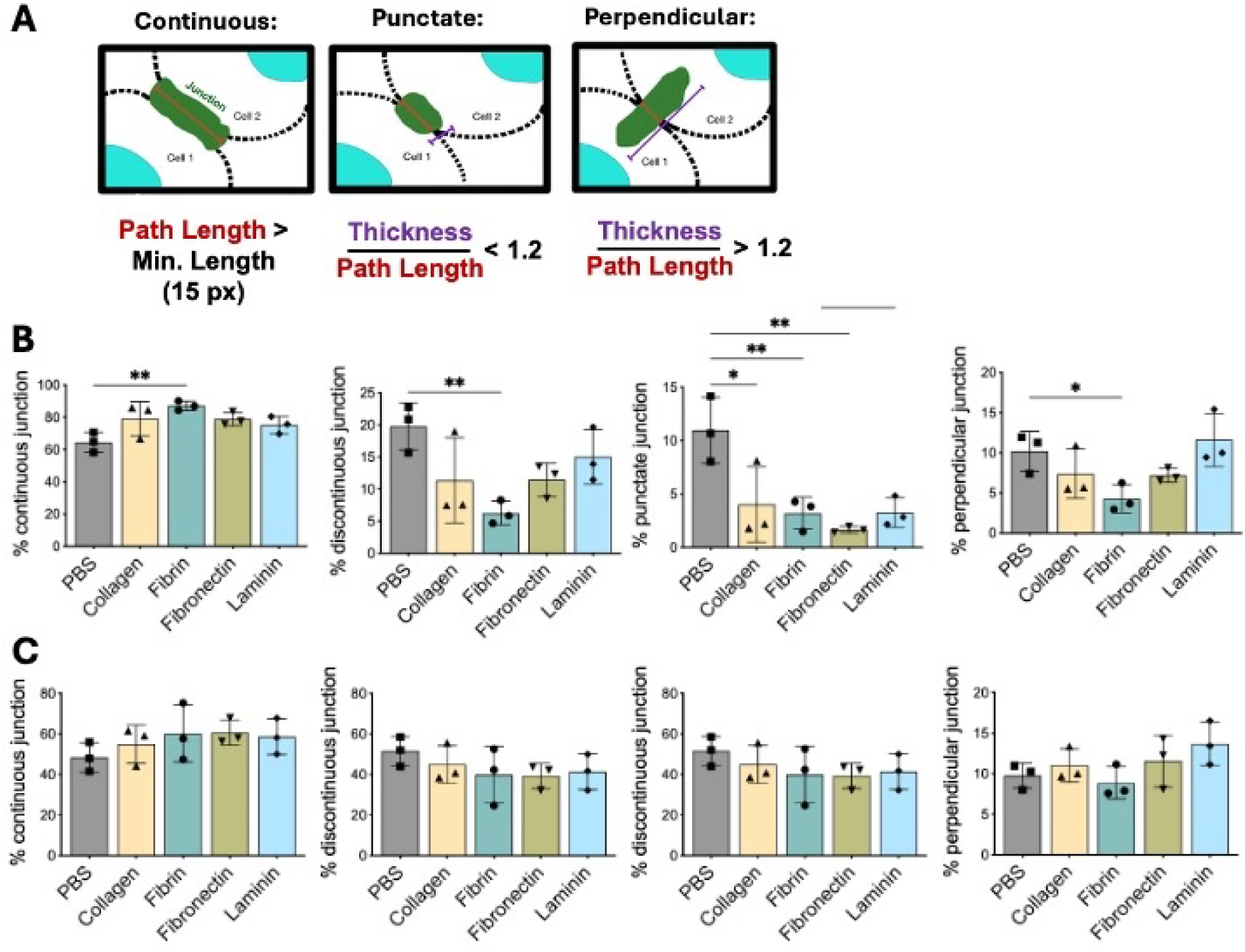
Fibrin coating enhances hLEC tight junction continuity. **(A)** Schematic of junction characterization using JAnaP. **(B)** Quantification analysis of ZO-1 junction morphology along cell perimeter on different ECM coatings. **(C )** Quantification analysis of VE-cadherin junction morphology along the cell perimeter on different ECM coatings. Data presented as mean ± SEM from n=3 experiments. Statistical significance was performed using ordinary One-way Anova with post hoc Dunnett test where significance levels are indicated as * p < 0.05, ** p < 0.01, *** p < 0.001, **** p < 0.0001; ns, not significant.

Fibrin yielded the most pronounced changes for ZO-1 with 87.1 ± 1.6 % continuous junctions and 0.06 ± 0.01 % discontinuous junctions compared to the PBS control (64.5 ± 3.5 and 0.19 ± 0.02 %, respectively). Collagen and fibronectin coatings had increased continuous junctions in ZO1, 79.0 ± 6.1 %, and 78.8 ± 2.3%, respectively. All ECM coatings substantially reduced punctate junctions, while only fibrin reduced perpendicular junctions by 0.04 ± 0.01%. Typically, perpendicular describes an elongated, cell-cell junction morphology potentially caused by cell-cell stretching or extension. This suggests that fibrin provides a stronger barrier, leading to less stretching or separation of junctions and, consequently, fewer perpendicular junction formations. In contrast, JAnaP analysis with VE-cadherin revealed no significant ECM-dependent differences in overall cell morphology or junctional continuity (**Figure 3C**).

### mRNA expression of lymphatic junction integrity-associated markers

We found that fibrin caused a reduction in RhoA gene expression, but no differences in junctional proteins, suggesting that translation/localization to the cell surface and not transcription are altered by ECM materials. qPCR analysis was performed to assess the expression of junctional protein genes and LEC marker genes. We found no significant differences in the expression of key adherens and tight junction components, including ZO-1, VE-cadherin, β-catenin, and claudin-5 for ECM proteins other than laminin. We found that laminin coated LECs have significant reduced of claudin-5 and increased VE-cadherin expression. LEC-specific transcription factor PROX1 and LYVE-1 remained unchanged across ECM coatings. Notably, fibrin coatings also led to a reduction in RhoA gene expression in LECs (**Figure 4**).

**Figure 4.**
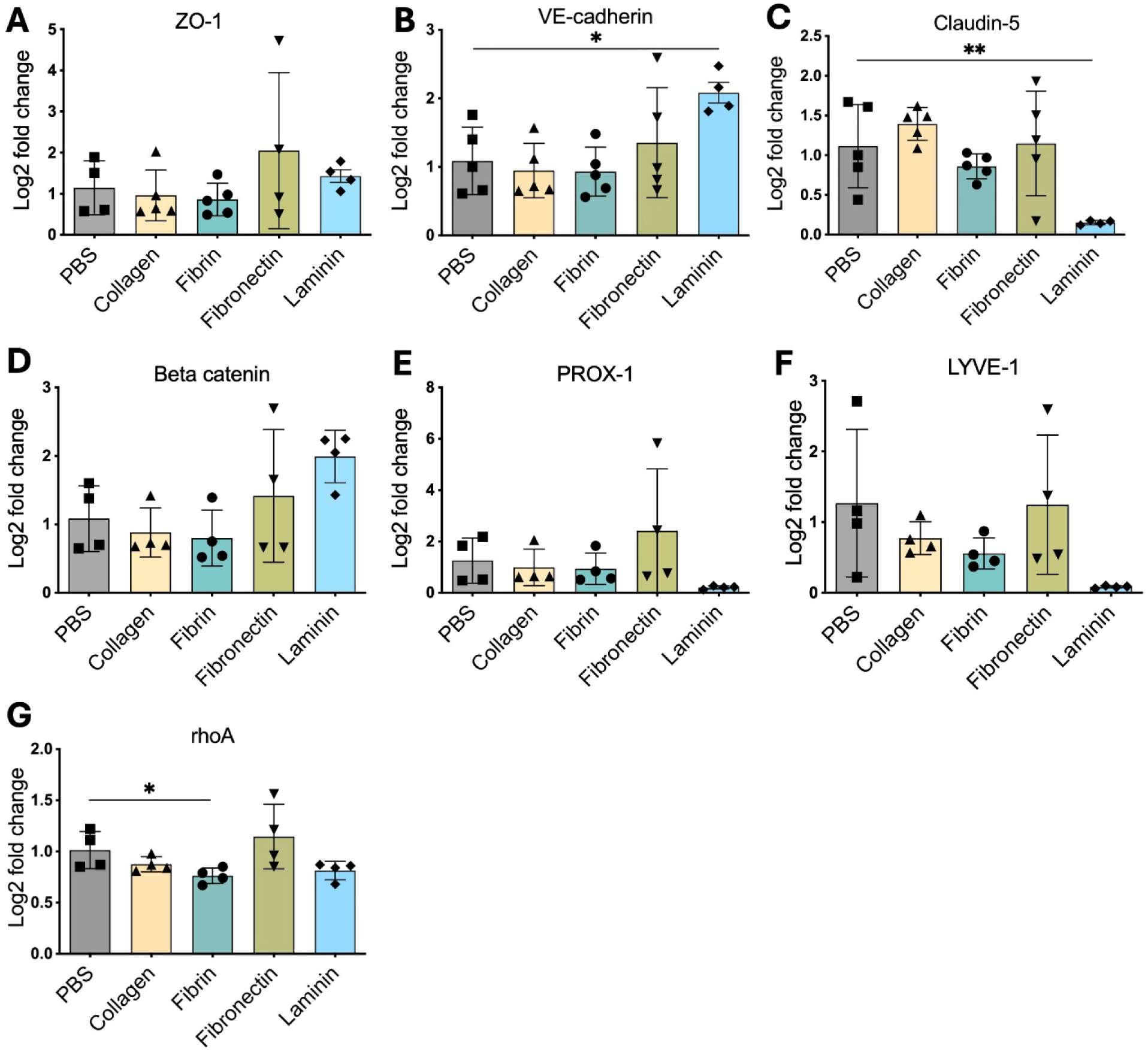
**Fibrin caused a significant difference in rho-A expression but we see no significant difference in relative gene expression of lymphatic junction integrity-associated proteins compared to no ECM control. (A) ZO-1 (gene: tight junction 1)**, (B) VE cadherin (gene: CDH5), (C ) Claudin 5 (gene: CLDN5), (D) β-catenin (gene: CTNNB1), (E ) PROX1, (F) LYVE-1, (G) rhoA. Data are presented as mean ± SEM from n=4 biological replicates RNA. Unpaired t-test was performed where significance levels are indicated as * p < 0.05, ** p < 0.01, *** p < 0.001, **** p < 0.0001; ns, not significant

## DISCUSSION

The goal of this study was to elucidate the relationship between ECM components and LEC junction morphology that determine vascular permeability. We evaluated the influence of collagen I, fibronectin, laminin, and fibrin coatings on LEC barrier integrity and solute transport. Culturing LECs on these ECM substrates over time revealed no significant differences in cell growth or density. Upon reaching confluence, LECs displayed the highest TEER values on fibrin, followed by collagen. Dextran transport assays corroborated these results as fibrin-coated LEC monolayers exhibited the lowest dextran permeability, consistent with enhanced barrier function. Supporting our findings, a recent study using HUVECs demonstrated that an aligned fibrin matrix strengthened endothelial junctions and increased barrier integrity compared to isotropic fibrin or fibronectin, with a 2.5-fold rise in relative expression of junction proteins, PECAM-1, occuldin, and ZO-1, and a 30% reduction in permeability ^24^. This shows that fibrin enhances endothelial barrier function. In contrast, studies on microvascular endothelial cells (MVECs) have reported opposite effects of fibrin(ogen). In human lung MVECs, the circulating fibrinogen degradation product fragment X induced endothelial dysfunction and disrupted barrier integrity of VE-cadherin ^25^. Likewise, in rat cardiac MVECs, fibrinogen exposure enhanced endothelial permeability via ERK pathway activation and stimulated F-actin formation ^26^. These findings suggest that fibrin(ogen) regulates endothelial permeability, potentially through cell- and tissue-type-specific mechanisms mediated between the cytoskeleton, focal adhesion proteins, and the ECM. These interactions ultimately modulate cell-cell junction expression, morphology, and permeability.

Extensive research has demonstrated that ECM composition governs cell function, differentiation, and physiological homeostasis, whereas abnormalities in ECM organization often contribute to disease. In the context of lymphatics, most ECM-related studies have focused on lymphangiogenesis and tube formation, where remodeling of ECM structure and associated proteins has been documented during development, disease progression, and wound healing. Lombardi *et al.* reported that cathepsin-L–deficient (nkt) mice exhibited contrasting alterations in ECM protein distribution where collagen, fibronectin, and laminin were upregulated in lymph nodes but downregulated in the thymus ^27^. Similarly, a single-cell RNA-seq cohort study identified COL1A1⁺ endothelial cells as contributors to tumor progression and metastasis in gastric cancer ^28^. These findings underscore the importance of defining ECM protein modules to better understand lymphatic vascularity. It is now recognized that lymphatic endothelial permeability is dynamically regulated by multiple factors including inflammatory mediators through interactions among endothelial, smooth muscle, and stromal cells. Our work showed that fibronectin coatings did not significantly alter LEC barrier function, while laminin reduced TEER and increased permeability. In a 3D lymphatic-on-a-chip model, Henderson *et al.* showed that incorporating fibronectin into a collagen I hydrogel enhanced junctional tightness via integrin α5-mediated signaling, thereby promoting vascular integrity ^29^. Immunofluorescence analysis in our study revealed that ZO-1 expression was significantly elevated on fibrin and collagen substrates, whereas the immunostaining of VE-cadherin expression (MFI) remained unchanged. JAnaP indicated that fibrin increased continuous ZO-1 junctions by ∼35% relative to control, while collagen and fibronectin showed ∼22% increases. Thus, our work corroborates the existing studies, suggesting that fibronectin and collagens increase junction continuity and thus increase the LEC barrier.

Our work has shown that individual ECM components, particularly fibrin and collagen, enhance LEC barrier function. Research has shown that, for other cell types, particularly those of the blood-brain barrier, ECM proteins promote barrier integrity. Human brain microvascular endothelial cells (HMBECs) cultured on fibronectin alone or a combination of fibronectin, collagen IV, and laminin exhibit more continuous VE-cadherin and ZO-1 junction morphology compared to other ECM coatings^30^. In addition, human umbilical vein endothelial cells (HUVECs) cultured on a 50% collagen I and 50% fibronectin coating demonstrated optimal cell spreading, contractility, and morphology compared to other collagen-to-fibronectin ratios ^31^. These suggest that ECM proteins work to regulate junction protein expression and barrier integrity is reflected across other endothelial cell types, not only LECs.

Our findings corroborate previous studies that found culturing LECs on ECM proteins has minimal or no consequential effect on the expression of key lymphatic markers. Gene expression profiling by qPCR showed no significant differences in key junctional genes (ZO-1, VE-cadherin, claudin-5, and β-catenin) or in the LEC-specific transcription factor PROX1 across ECM coatings for ECM substrates other than laminin. This suggests that ECM proteins, especially fibrin and collagen I, may not directly affect junctional protein expression at the gene level but instead regulate their trafficking and organization locally. A decrease in claudin-5 expression only in laminin could be the reason for high LEC permeability with the laminin substrate. Previous work by Frye *et al.* demonstrated that LECs cultured on soft matrices upregulated genes related to cell migration and lymphangiogenesis^32^, and Saha *et al.* reported that dopamine-conjugated hyaluronic acid coatings preserved PROX1 and LYVE-1 expression more effectively than fibronectin^33^. In our study, fibrin coating decreased RhoA gene expression, suggesting reduced stress fiber formation compared to other ECMs; however, anisotropy analysis indicated no major difference in F-actin organization among groups. Taken together, these findings suggest that individual ECM proteins do not significantly compromise the preservation of key lymphatic markers or the alignment of the actomyosin fibers making up the cytoskeleton=.

## Conclusion

Our findings demonstrate that ECM composition critically influences LEC junctional integrity and permeability. Among the tested matrices, fibrin and collagen most effectively promoted barrier function, while fibronectin and laminin exhibited lesser or opposite effects. These results provide new insights into how ECM composition modulates lymphatic endothelial barrier properties and may guide the design of physiologically relevant ECM matrices for modeling lymphatic permeability *in vitro*. Studies using these more physiologically relevant models may then shed light on how, in the context of the disease and inflammation, changes in the extracellular environment may modulate lymphatic barrier functions.

## Acknowledgements

We would like to thank the UMD Bioworkshop core facility and Dr. Kimberly Stroka’s Cell and Microenvironment Engineering Lab, specifically Ken Brandon, for assistance and input with the Junction Analyzer Program. This work was funded by NIH MIRA R35 (KM, SAE, AA), the NIH Diversity Supplement (AA), and the Drs. Wayne T. & Mary T. Hockmeyer Summer Fellowship (AA). We’d also like to acknowledge claude.ai for aid in editing the abstract of the manuscript.

## MATERIALS AND METHODS

### Cell Culture and Preparation of Extracellular Matrix Coating

Primary human dermal lymphatic endothelial cells (LECs) (Promocell, C12217) were cultured on rat tail collagen type I in Endothelial Cell Growth Microvascular Media (EGMV2) (Promocell, C-22121) supplemented with insulin-like growth factors, fibroblast growth factors with fetal calf serum. We used three batches of LECs (487Z030.1, 495Z030.4, 521Z007.2), which were derived from females aged 42-60 years, and expanded them for our cell stocks.

12-well transwells and 8-well glass-bottom chamber slides were coated with a final concentration of 50µg/mL of the following ECM proteins prior to seeding LECs: Collagen, Fibrin, Fibronectin and Laminin. Collagen coating solution composed of rat tail collagen type I (Corning 354236), 1% acetic acid, and PBS. Fibronectin (MiliporeSigma FC010) and laminin (Sigma-Aldrich L4544) from human plasma were reconstituted in PBS. To prepare fibrin, fibrinogen from human plasma (Sigma-Aldrich F3879) was reconstituted in PBS and mixed with 4U of thrombin.

### Dextran Permeability Assay

The apical side or bottom of 1.0 µm pore size, 12 mm transwell inserts (Falcon, 353103) were used. Transwell membranes were coated with either collagen, fibrin, fibronectin, and laminin. LECs were seeded at 250,000/cm^2^ density and allowed to grow until confluence. Transendothelial electrical resistance (TEER) was measured by placing an electrode in the top and bottom wells (basolateral and apical side). 4kDa FITC-dextran was used to measure transport. Dextran was added to the top compartment, basolateral side. Bottom well, apical side was sampled up to 8 hours.

The overall TEER of the monolayer (ohms.cm^2^) was determined by multiplying the TEER values by the area of the monolayer on transwell. Transport was quantified by correlating the fluorescence intensity of dextran in the bottom well with the mass of dextran.

### Immunofluorescence Imaging

Transwells were fixed with 2% Paraformaldehyde and permeabilized with Triton-X100. Cells were blocked with 2% fetal bovine serum (FBS) followed by cut with dissection scissors and moved to glass slides. LECs are immunostained with primary antibodies human anti-mouse VE-cadherin (BD Biosciences, 55566) and rabbit anti-human ZO-1 (Thermofisher, 33-9100) at 4^0^C overnight in 2% FBS and stained with secondary antibodies donkey anti-mouse (Invitrogen, A-21202, A-31571 and donkey anti-rabbit (Invitrogen, A-21206, A-31573) in addition to phalloidin (Invitrogen A12379, A34055, A22287) for 2 hours at room temperature in 2% FBS. Following both primary and secondary antibodies, LECs were washed with 0.1% Tween-20. LECs were stain for their nuclei with Hoescht nuclei stain for 10 minutes and then mounted with Vectashield mounting media.

### Image Analysis

To conduct post-imaging analysis, Image J/FIJI and the Junction Analyzer Program (JAnaP) was used. Using the Cell Counter Plugin with FIJI, the nuclei of the LEC monolayer were counted and scaled to the monolayer area to determine total cell count. JAnaP was used to characterize junction morphology as continuous or discontinuous (Gray, K et al 2019 & 2020). Discontinuous was further characterized as perpendicular or punctate junction morphology. Additional morphological features, including circularity, solidity, and area was also determined with JAnaP.

### RNA isolation and qPCR

To quantify relative gene expression following LECs being cultured on different ECM compositions, LECs’ RNA was isolated using the TRIzol (Invitrogen, 15596026) isolation method. 3-4 wells were pooled to collect sufficient RNA. RNA was reverse transcribed to create complementary DNA (cDNA) using a cDNA reverse transcription kit (Applied Biosystems, 4374966). TaqMan Universal PCR Master Mix (Applied Biosystems, 4444556) and PCR Probe and Primers detailed in Table 1 were used to conduct qPCR analysis.

**Table 1:**
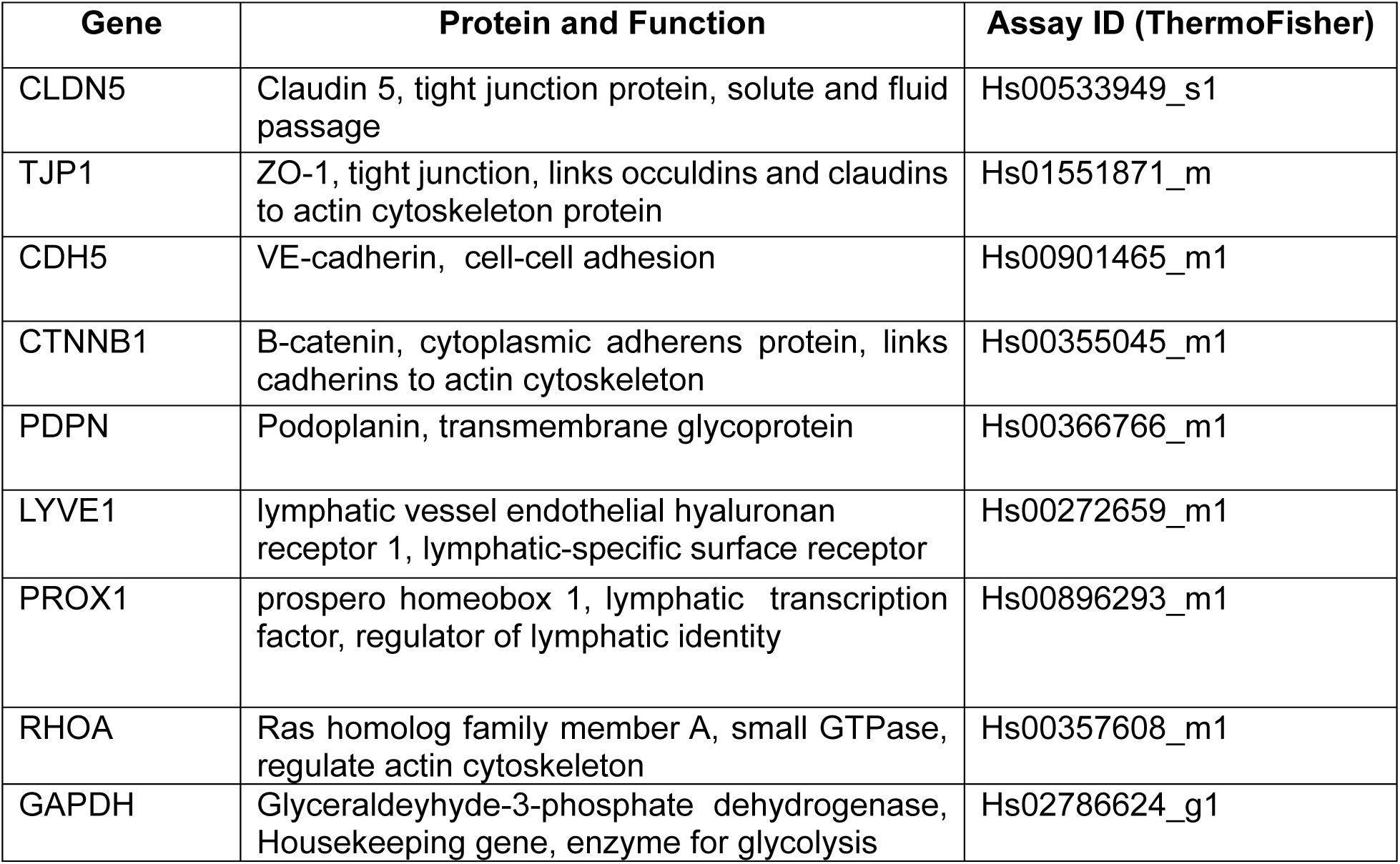
PCR Probes and Primers.

### Statistical Analysis

Group Analysis was performed for more than one variables using a two-way ANOVA and one-way ANOVA and Welch’s test were done with multiple comparison for single factor group comparison. Statistical significance will be established at p<0.05, and all analyses will be performed using GraphPad software v10. Results will be expressed as mean ± standard error of the mean (SEM).

## Notes

### Competing Interest Statement

The authors have declared no competing interest.

