## Supplementary figures and images for "Extracellular matrix proteins modulate lymphatic endothelial cell junction morphology and barrier function"

### Supplemental Figure

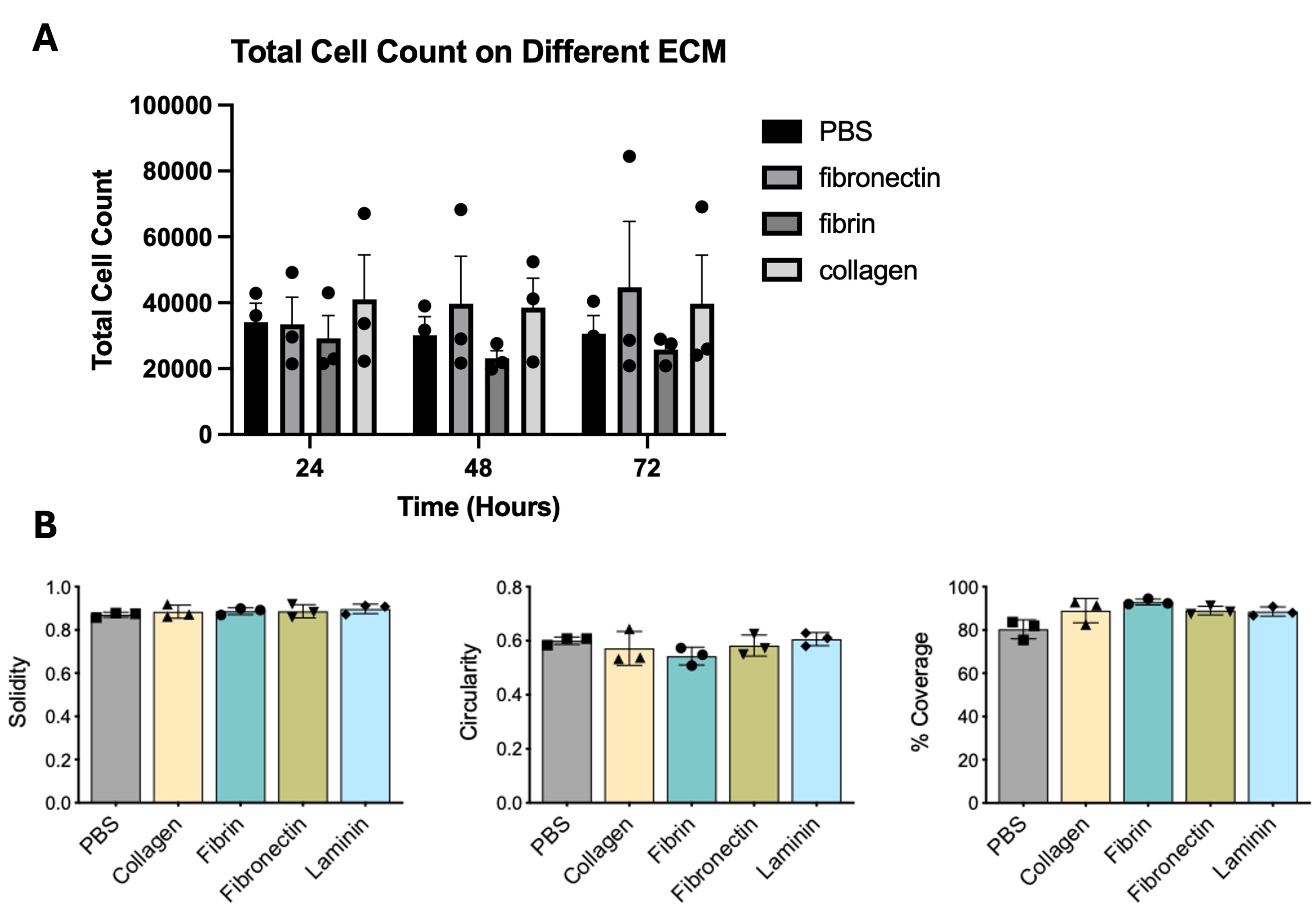
